# Health Status by Gender, Hair Color, and Eye Color: Red-Haired Women are the Most Divergent with the Lowest Viability and the Highest Fertility

**DOI:** 10.1101/139295

**Authors:** P. Frost, K. Kleisner, J. Flegr

## Abstract

**Background:** Red hair is associated with pain sensitivity, and more so in women than in men. Hair redness may thus interact with a female-specific factor. We tested this hypothesis on a large sample of Czech and Slovak respondents. They were asked about the natural redness and darkness of their hair, their natural eye color, their physical and mental health (24 categories), and other personal attributes (height, weight, number of children, lifelong number of sexual partners, frequency of smoking).

**Results:** We found that red-haired women did worse than other women in ten health categories and better in only three. In particular, they were more prone to colorectal, cervical, uterine, and ovarian cancer. Cancer risk increased steadily with increasing hair redness except for the reddest shade. Red-haired men showed a balanced pattern of health effects, doing better than other men in three categories and worse in three. Number of children was the only category where both male and female redheads did better than other respondents. We also confirmed earlier findings that red hair is naturally more frequent in women than in men.

**Conclusion:** Red-haired women had higher fecundity and sexual attractiveness, but this selective advantage seems offset by worse health outcomes and therefore lower viability. The resulting equilibrium between these two counterbalancing forces might explain why red hair has remained less common than other hair and eye colors. Of the ‘new’ hair and eye colors, red hair diverges the most from the ancestral state of black hair and brown eyes. It is the most sexually dimorphic variant, not only in population frequency but also in health outcomes. This sexual dimorphism seems to have resulted from a selection pressure that acted primarily on early European women and which led to a general and apparently rapid diversification of hair and eye colors.

## Background

It has long been known that redheads are at higher risk of sunburn and skin cancer. This is to be expected because red hair is associated with fair skin, which is more vulnerable to the harmful effects of UV radiation [1]. Less expectedly, red hair is also associated with increased pain sensitivity and a higher risk of endometriosis and Parkinson’s disease [2,3,4,5,6,7]. These associations seem to involve a risk factor not directly related to fairness of skin and vulnerability to UV.

This risk factor seems to be specific to women. Pain sensitivity is greater in female redheads than in male redheads, and the association between red hair and endometriosis is obviously specific to women [2,5]. Red hair alleles promote Parkinson’s disease by compromising the integrity of dopaminergic neurons, but no one has determined whether this disorder affects red-haired women more often than red-haired men [8]. If a female-specific factor is interacting with red hair to facilitate these medical conditions, a plausible candidate may be higher levels of estrogen in the fetal environment, which promote not only certain health outcomes but also certain hair and eye colors. Prenatal estrogen particularly seems to favor red hair. According to a twin study, women are likelier than men to have red hair even when the genotype is the same [9]. Prenatal estrogen may also affect eye color. Face shape is more feminized in blue-eyed men than in brown-eyed men of the same ethnic background [10,11]. In addition to favoring blue eyes over brown eyes, prenatal estrogen seems to favor green eyes over blue eyes, the so-called blue-eye genotype being expressed in women more often as green eyes [12]. In sum, there seems to be a general tendency for women to exhibit less frequent hair and eye colors at the expense of more frequent ones [13,14].

Prenatal estrogen may therefore mediate the apparent effect of red hair on certain health outcomes, including some that remain unsuspected. It was only by chance that researchers discovered the three-way association between being a woman, having red hair, and feeling more sensitivity to pain. There has been no systematic effort to identify all female-specific associations between human health and red hair, let alone between human health and each of the different hair and eye colors.

For these reasons, we wished to find out how different aspects of human health vary as a function of hair/eye color. We also wished to see how well the variance is explained by the two known risk factors: 1) vulnerability to UV, as measured by relative importance of skin cancer; and 2) gender, specifically being a woman. To this end, we used an existing pool of data collected for an unrelated purpose: a survey on the effects of RhD factor on various health categories in a Czech and Slovak population. This survey encompassed a very large number of individuals and could thus capture small effects. On the other hand, the data were collected by self-report, a method prone to noise because of differences in self-evaluation among respondents.

## Methods

### Respondents and recruitment

The present study reanalyzed data originally collected for a survey on the effects of RhD factor on human health. Respondents were recruited by a Facebook-based snowball method [15], as described by [16]. In short, potential volunteers were invited to participate in “research to investigate how blood groups and other biological factors relate to personality, performance, morphology, and health.” The invitation was posted on the Facebook wall page “Guinea Pigs” (in Czech “Pokusni kralici”) for Czech and Slovak nationals willing to take part in evolutionary psychology experiments (www.facebook.com/pokusnikralici) [17]. The first page of the electronic questionnaire described the goals of the study. The following note was also included: “The questionnaire is anonymous and obtained data will be used exclusively for scientific purposes. Your cooperation with the project is voluntary, and you can terminate it at any time by exiting this website.” The first and final pages of the questionnaire both had a Facebook share button and the following request: “We need the highest possible number of respondents. Therefore, please share the link to this questionnaire with your friends, for example on Facebook.” The share button was pressed by 575 respondents, with the result that we finally obtained data from 7,044 Czechs and Slovaks between 28/4/2014 and 12/09/2016. The study, including the method of obtaining informed consent (by pressing the Next button on the first page), was approved by the IRB of the Faculty of Science, No. 2014/21.

### Questionnaire

The questionnaire was distributed as a Czech/English Qualtrics survey (http://1url.cz/q05K). In the first part of the questionnaire, respondents were asked, among other questions, to rate the natural darkness of their hair and eyes and the natural redness and waviness/curliness of their hair on a 6-point Likert scale (0- light, not curly/wavy/red at all 5- very dark, red, curly/wavy). They were also asked to choose the natural color of their eyes from a list of eight colors (blue, green, brown, black, grey, amber, hazel, yellow). Finally, they were asked about their body height and weight, number of children, lifelong number of sexual partners, and how often they smoked (0- never, 1- maximum of once per month, 2-maximum of once per week, 3- maximum of once per day, 4- several times a day, 5- more than 20 cigarettes a day, 6- more than 40 cigarettes a day).

The medical part of the questionnaire was prepared by two physicians: a clinician (internist/hematologist), and a researcher (molecular geneticist). Questions fell into two parts, one using subjective measures of health status and the other more objective measures. Respondents were first asked to rate the presence and intensity of their health problems on a 6-point Likert scale. These questions were on physical health and mental health in general, and on more specific health categories: allergies; cancer; digestion; fertility; genitourinary system; heart and vascular system; hematology; immune system; metabolism, including endocrine system; musculoskeletal system; nervous system; respiratory organs; sense organs; and sexual function. The second part of the questionnaire was designed to provide objective information on health status. For example, respondents were asked how many physician-prescribed drugs they were currently taking per day, how many “different herbs, food supplements, multivitamins, superfoods etc.” they were currently taking per day, and how often they had used antibiotics during the past 365 days.

As a benchmark for the relative importance of associations between hair/eye color and the 24 health categories, we looked for associations between these categories and two unrelated but well-known risk factors: body mass index (BMI) and smoking. Finally, as another benchmark, we looked for significant associations between hair waviness and these categories.

For some of these categories, we also asked the respondents to state the specific disorders they had or used to have. For the ‘Cancer’ category, respondents were asked “What kind of cancer are you suffering from or have you suffered from?” They then read a list of disorders and ticked the appropriate boxes.

### Data analysis

Statistica v. 10 and IBM SPSS v. 21 were used for most of the statistical analysis. MANCOVAs on health effects (with gender, eye color, or hair color as predictors) were performed by “adonis” function available within Vegan package in R [18]. Differences in age were tested by t-tests. Chi^2^ tests were used to compare the frequencies of eye colors in men and women. Effects of gender and age on hair color, eye color, and waviness of hair were analyzed by both nonparametric (partial Kendall correlation with age or gender as a covariate) and parametric tests using general linear models with gender, age, and gender-age interaction as independent variables. Both categories of tests produced very similar results and therefore only the results of the more conservative nonparametric tests are reported. Logistic regression (Quasi-Newton estimation method) was used for the analysis of relation between cancer and hair redness and hair redness*gender interaction. Other ordinal and binary data were analyzed by a partial Kendall correlation test, which is used to measure the strength and significance of associations between binary, ordinal, or continuous data regardless of their distributions and which can be used to control for one confounding variable, here the respondent’s age [19,20]. To compute partial Kendall Tau and the significance of each variable, after controlling for age, we used an Excel spreadsheet available at: http://web.natur.cuni.cz/flegr/programy.php. Because many disorders tend to be more common in men than in women or vice versa, associations between the health categories and the hair/eye colors were always analyzed separately for men and women. Whenever fewer than 10 respondents reported a disorder, a Fisher exact test was used to determine the significance of the association between hair/eye color and the category of fitness. To correct for multiple tests, and the associated increase in the false discovery rate, we used the Benjamini-Hochberg procedure with the false discovery rate preset to 0.25 [21]. In contrast to the Bonferroni correction, this procedure takes into account the distribution of p values of performed multiple tests. Therefore, when the studied factor has multiple effects, the number of significant results after the correction may be higher than before the correction. To measure eye color diversity, we used the Simpson index λ, which was computed as λ = SUMA(p_i_)^2^, where p_1_–p_i_ denotes proportions of respondents with 1-i eye color in the population under study [22]. This index reflects the probability of two randomly-selected respondents carrying the same character, here the same eye color. All raw data are available as Supporting Information S1, at https://figshare.com/s/6a02dd5cec0f90b69db9

## Results

### Characteristics of respondents

Information on eye color, hair color, and hair waviness was provided by 2,558 men and 4,472 women out of 7,030 Czech and Slovak respondents (the others did not complete the questionnaire part of the test). Mean age of the men (36.8, std. dev. 13.5) was somewhat higher than mean age of the women (34.6, std. dev. 13.0) t_7028_ = 6.9, p < 0.0005.

Figures 1, 2, and 3 show how different gradations of hair redness, hair darkness, and eye darkness were distributed among male and female respondents. In keeping with the findings of an earlier twin study, red hair was more frequent in women than in men [9]. Hair tended to be lighter in women than in men, but eyes were equally dark. A closer look at the data, however, showed that eye color was more diverse in women than in men, with green eyes being more frequent in women and blue and brown eyes more frequent in men (Figure 4 and Table 1). This gender difference in eye-color diversity is seen in a higher Simpson index for men (0.263) than for women (0.234). Women had higher eye-color diversity in all 5-year age groups, except for the 41-45 age group.

Age was associated in women with darker eyes, darker hair, redder hair, and less wavy hair (Table 2), and in men with redder hair and less wavy hair. After controlling for the effect of age, we still found gender differences: male hair was significantly darker (mean 4.07 vs. 3.87, p < 0.0005), less red (mean 2.09 vs. 2.51, p < 0.0005), and less wavy (mean 2.30 vs. 2.56, p < 0.0005). No gender difference was observed in eye darkness (mean 3.39 vs, 3.42, p = 0.56).

**Figure 1.**
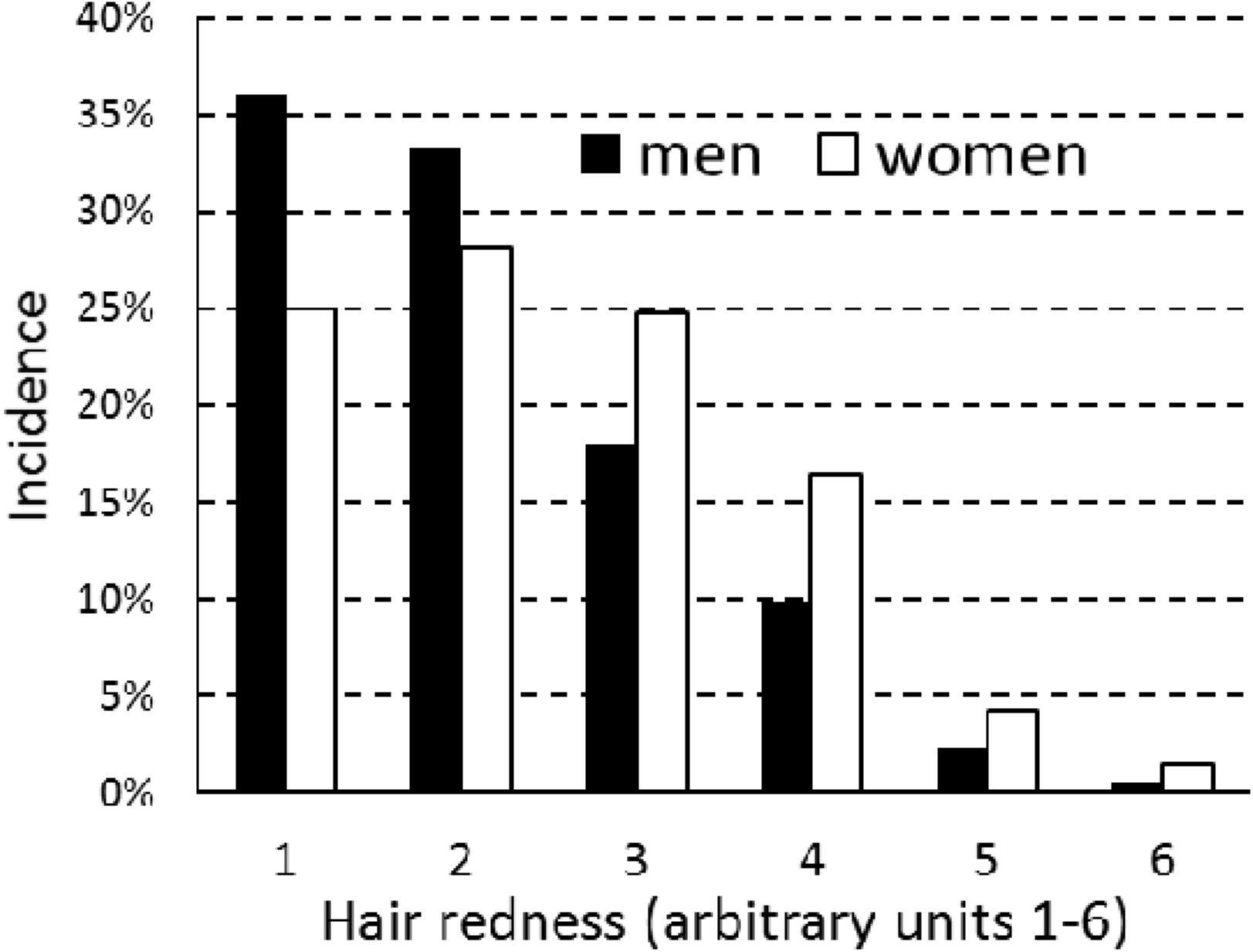
Gradations of hair redness: population frequencies for men and women

*Respondents rated hair redness on a scale of 1 to 6 where 1 = not at all red and 6 = completely red*

**Figure 2.**
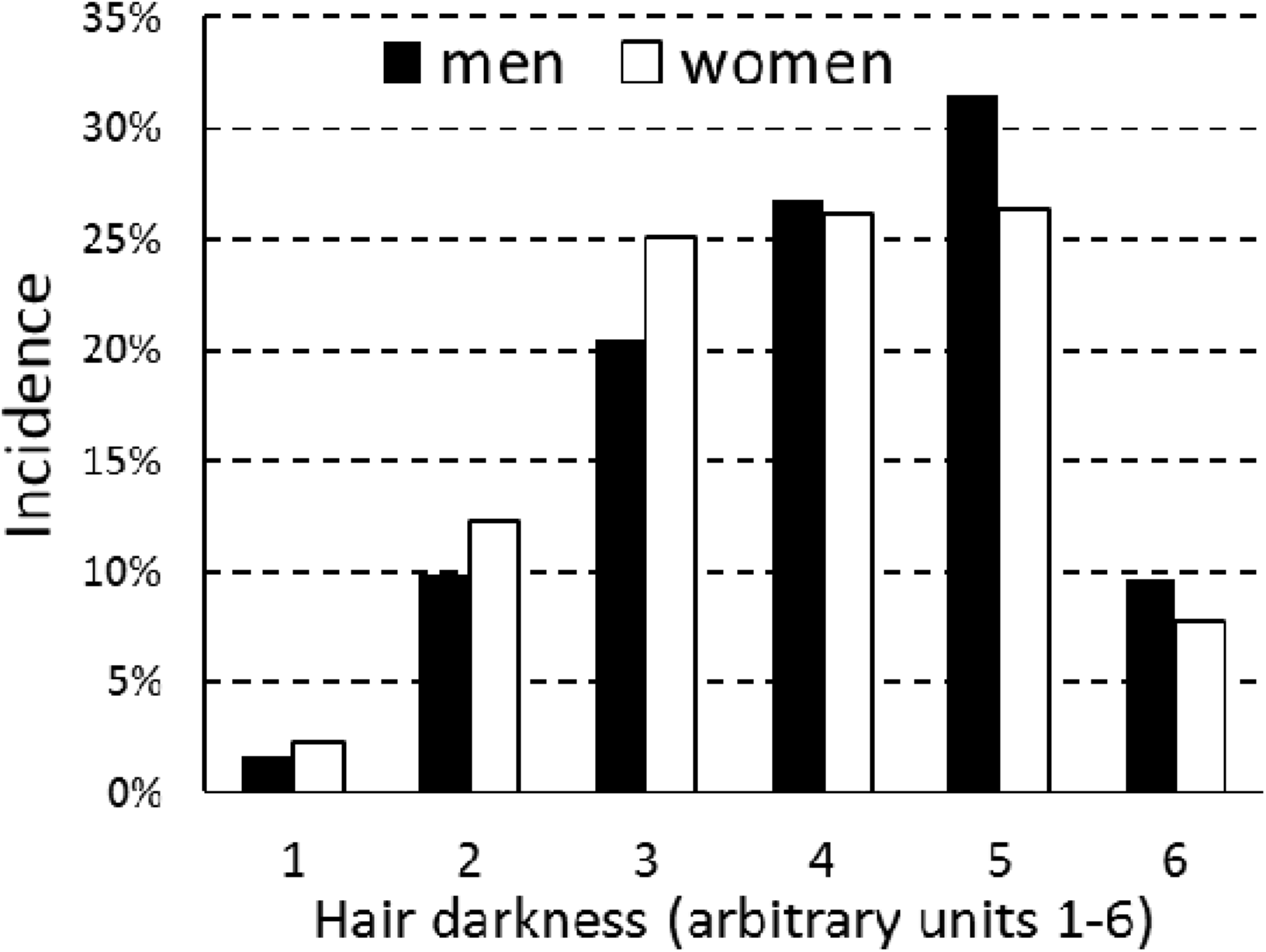
Gradations of hair darkness: population frequencies for men and women

*Respondents rated hair darkness on a scale of 1 to 6 where 1 = not at all dark and 6 = completely dark*

**Figure 3.**
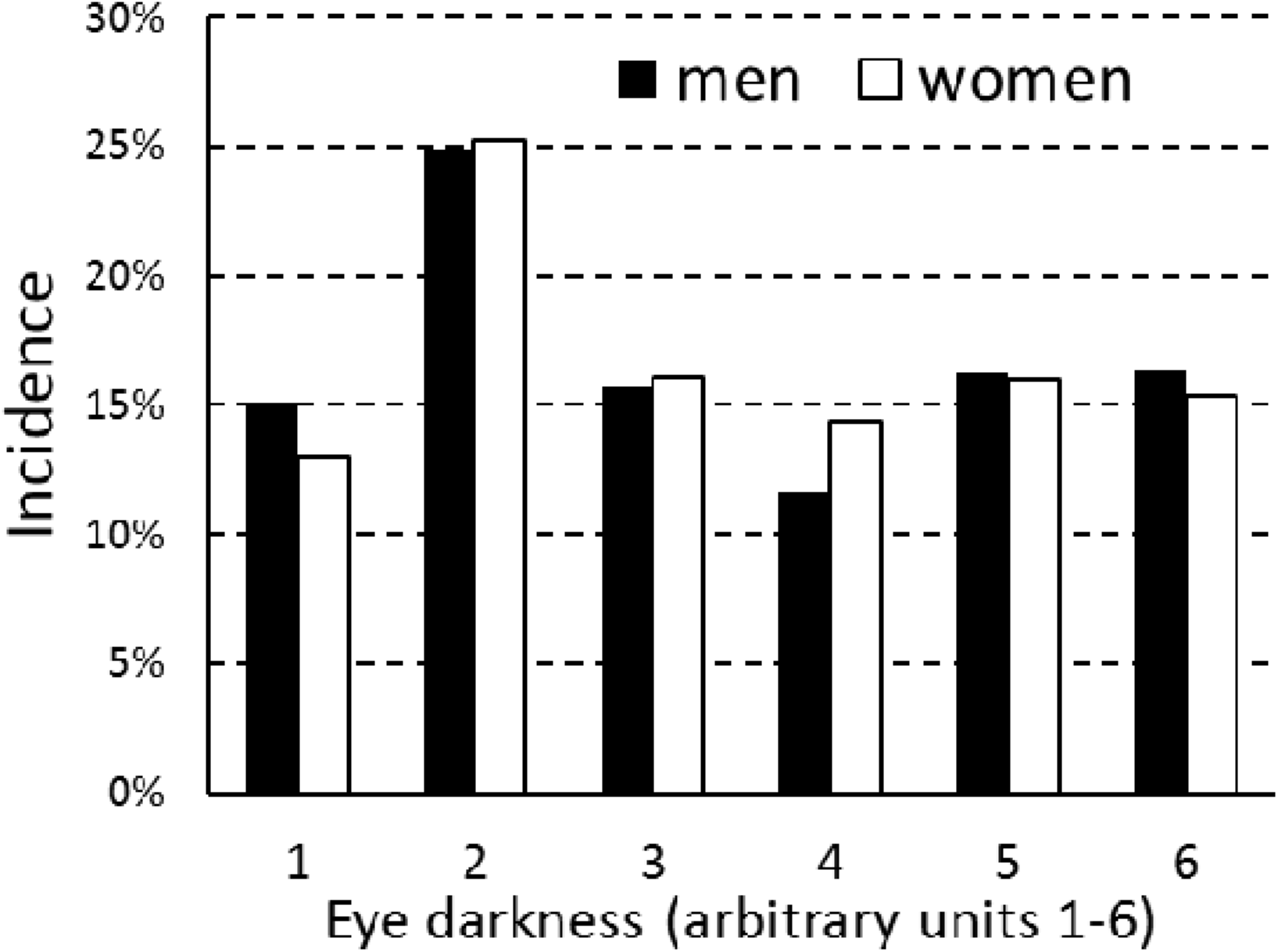
Gradations of eye darkness: population frequencies for men and women

*Respondents rated eye darkness on a scale of 1 to 6 where 1 = not at all dark and 6 = completely dark*

**Figure 4.**
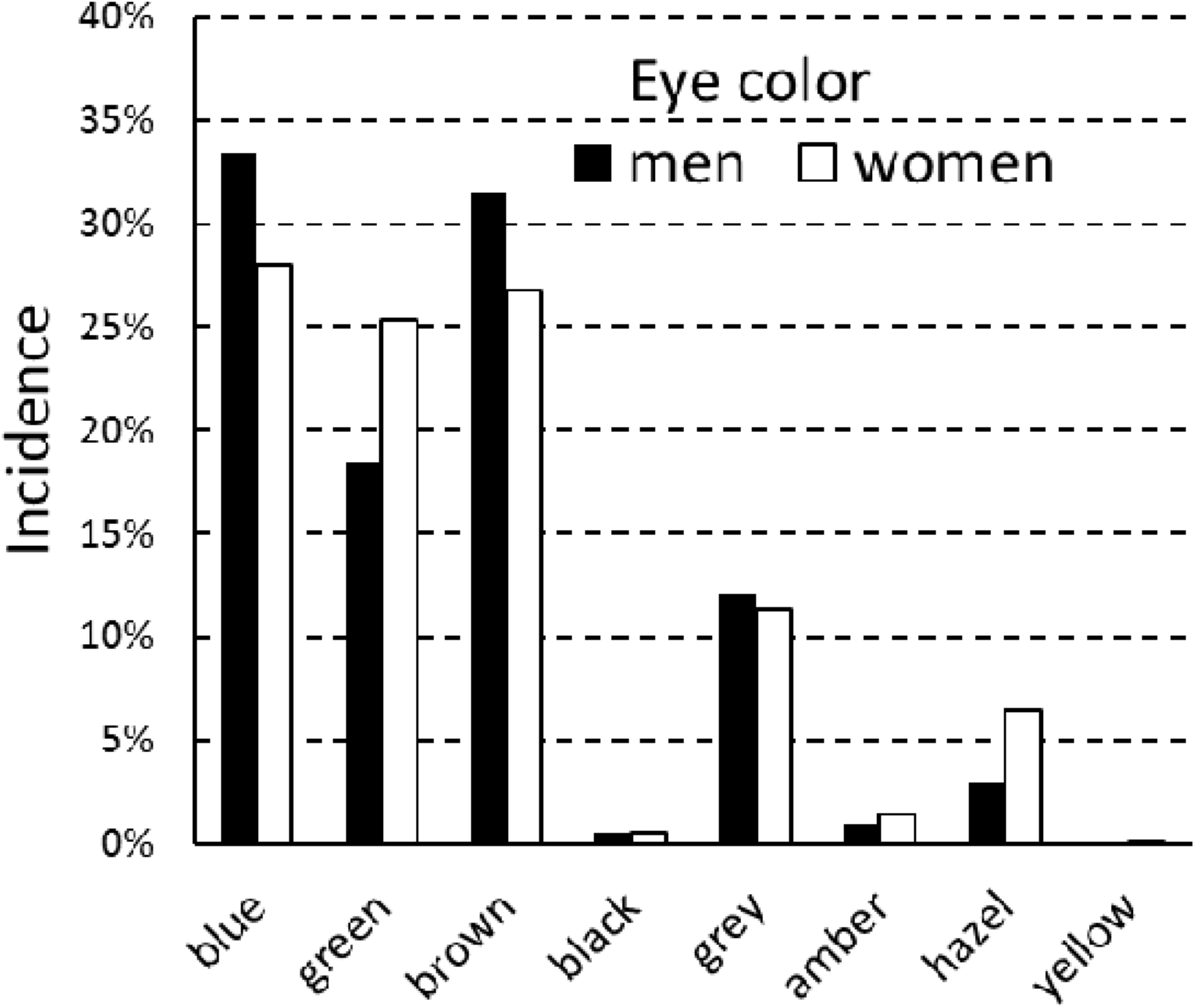
Eye colors: population frequencies for men and women

**Table 1.** Frequencies of eye colors in men and women

|  |  | Blue | Green | Brown | Black | Gray | Amber | Hazel | Yellow | Totals |
| --- | --- | --- | --- | --- | --- | --- | --- | --- | --- | --- |
| N | Men | 846 | 453 | 813 | 14 | 312 | 21 | 72 | 0 | 2531 |
| Percent |  | 33.43% | 17.90% | 32.12% | 0.55% | 12.33% | 0.83% | 2.84% | 0.00% | 100% |
| N | Women | 1243 | 1135 | 1203 | 19 | 513 | 66 | 258 | 5 | 4442 |
| Percent |  | 27.98% | 25.55% | 27.08% | 0.43% | 11.55% | 1.49% | 5.81% | 0.11% | 100% |
| N | All | 2089 | 1588 | 2016 | 33 | 825 | 87 | 330 | 5 | 6973 |
| Chi2 |  | 22.7 | 53.5 | 19.9 | 0.54 | 0.94 | 5.63 | 31.0 | 2.85 |  |
| p |  | 0.000 | 0.000 | 0.000 | 0.463 | 0.333 | 0.018 | 0.000 | 0.091 |  |

*Benjamini-Hochberg correction for multiple (8) Chi^2^ tests indicated that gender differences were significant for all eye colors, except for black, gray, and yellow. The p values lower than 0.0005 were coded as 0.000.*

**Table 2.** Correlations between eye/hair properties and age

|  | Hair<br>darkness | Hair<br>redness | Hair<br>waviness | Eye<br>darkness | Age<br>(men) | Age<br>(women) |
| --- | --- | --- | --- | --- | --- | --- |
| Hair darkness |  | **-0.18** | **0.05** | **0.38** | 0.00 | **0.07** |
| Hair redness | -0.01 |  | **0.08** | **-0.08** | **0.14** | **0.18** |
| Hair waviness | **0.09** | **0.07** |  | 0.00 | **-0.09** | **-0.06** |
| Eye darkness | **0.39** | 0.00 | 0.00 |  | 0.02 | **0.03** |

*The upper-right part of the table (excluding the last two columns) shows the partial Kendall Tau correlations (age controlled) for men, and the lower-left the same results for women. The last two columns show standard Kendall Tau correlations between hair/eye properties and age in men and women, respectively. Significant correlations are in bold.*

### Associations between health categories and hair/eye color

We looked for significant associations between 24 health categories and different hair or eye colors. The results are shown in Table 3 for men and in Table 4 for women. Yellow eye color was reported by only 5 respondents and therefore excluded from the analyses.

**Table 3.** Results for men: associations with eye color, hair color, hair waviness, BMI, and smoking

|  | Blue | Green | Brown | Black | Gray | Amber | Hazel | Eye<br>darkness | Hair<br>darkness | Hair<br>redness | Hair<br>waviness | BMI | Smoking |
| --- | --- | --- | --- | --- | --- | --- | --- | --- | --- | --- | --- | --- | --- |
| Physical health<br>problems in<br>general | -0.015 | -0.019 | 0.011 | **0.033** | **0.041** | **-0.030** | **-0.025** | 0.004 | 0.015 | **-0.031** | **0.025** | **0.172** | **0.093** |
| Mental health<br>problems in<br>general | **-0.026** | 0.009 | 0.014 | 0.014 | 0.015 | 0.003 | **-0.024** | 0.011 | -0.024 | **0.032** | 0.014 | **-0.023** | **0.083** |
| Specific health<br>problems: |  |  |  |  |  |  |  |  |  |  |  |  |  |
| Antibiotics/<br>year | 0.007 | -0.001 | -0.004 | **-0.032** | 0.002 | **-0.036** | **0.026** | -0.018 | -0.002 | 0.006 | **0.025** | 0.014 | **-0.050** |
| Acute care/<br>5 years | 0.008 | **-0.034** | **0.048** | -0.006 | **-0.034** | -0.004 | -0.007 | 0.026 | 0.014 | -0.008 | **0.045** | 0.011 | **-0.034** |
| Different med.<br>specialist seen<br>every 2 years | 0.002 | -0.003 | 0.002 | 0.026 | 0.002 | -0.015 | -0.012 | -0.001 | -0.021 | 0.017 | **0.038** | **0.083** | **-0.030** |
| Number of<br>prescription | 0.015 | 0.023 | -0.014 | **0.051** | -0.017 | -0.012 | **-0.039** | -0.016 | -0.026 | **0.031** | 0.009 | **0.133** | **-0.053** |
| drugs/day |  |  |  |  |  |  |  |  |  |  |  |  |  |
| Number of<br>alternative<br>medicines/day | 0.021 | 0.000 | -0.019 | -0.020 | 0.001 | -0.015 | 0.007 | -0.003 | -0.010 | **0.032** | **0.041** | 0.013 | **-0.049** |
| Allergies | -0.001 | 0.003 | -0.004 | -0.014 | 0.015 | 0.020 | **-0.028** | -0.012 | 0.015 | **-0.053** | 0.016 | -0.001 | **-0.037** |
| Immunological | **0.024** | **-0.051** | 0.019 | -0.003 | 0.023 | -0.023 | **-0.037** | -0.005 | -0.003 | 0.000 | **0.029** | -0.015 | **-0.034** |
| Digestive | **-0.033** | 0.018 | 0.004 | 0.025 | **0.034** | 0.001 | **-0.040** | 0.020 | -0.017 | 0.012 | 0.013 | **0.023** | **0.053** |
| Heart &<br>vascular<br>system | -0.022 | 0.012 | -0.007 | 0.003 | **0.037** | **0.040** | **-0.044** | 0.003 | 0.005 | -0.008 | **0.026** | **0.076** | **0.025** |
| Hematological | 0.018 | 0.018 | -0.015 | -0.023 | 0.003 | -0.011 | **-0.042** | -0.028 | -0.017 | 0.024 | 0.019 | -0.002 | -0.009 |
| Metabolic | 0.018 | -0.013 | -0.001 | 0.013 | 0.012 | -0.009 | **-0.042** | -0.007 | -0.024 | 0.022 | 0.020 | **0.163** | -0.017 |
| Cancer | 0.013 | -0.030 | -0.015 | -0.010 | **0.026** | **0.053** | -0.004 | 0.000 | -0.020 | 0.018 | **0.029** | **0.039** | **-0.019** |
| Fertility | -0.015 | -0.023 | 0.012 | **0.074** | **0.028** | 0.012 | **-0.036** | 0.006 | 0.000 | -0.003 | -0.001 | 0.017 | 0.001 |
| Genitourinary | 0.016 | -0.008 | -0.004 | -0.024 | -0.009 | 0.001 | 0.013 | -0.013 | -0.025 | 0.018 | 0.004 | -0.011 | -0.028 |
| Sense organs | **-0.034** | -0.002 | 0.025 | -0.005 | **0.027** | -0.004 | -0.018 | 0.023 | 0.005 | 0.004 | 0.013 | 0.017 | 0.009 |
| Neurological | **-0.028** | -0.023 | 0.007 | 0.003 | **0.059** | -0.012 | -0.001 | 0.008 | 0.005 | 0.023 | 0.018 | -0.018 | -0.012 |
| Psychiatric | **-0.037** | 0.001 | 0.014 | **0.042** | 0.007 | 0.002 | **0.030** | 0.027 | 0.008 | 0.008 | **0.038** | -0.009 | **0.095** |
| Sexual<br>function | -0.003 | 0.003 | -0.011 | 0.020 | **0.029** | -0.006 | **-0.032** | -0.022 | -0.011 | 0.007 | -0.003 | **0.034** | **0.074** |
| Musculoskeletal | **-0.052** | -0.025 | **0.063** | 0.014 | 0.016 | 0.010 | -0.017 | **0.066** | 0.001 | 0.026 | **0.034** | **0.037** | **-0.021** |
| Respiratory | -0.011 | -0.013 | 0.030 | 0.019 | -0.006 | 0.010 | **-0.027** | 0.014 | 0.006 | 0.001 | 0.017 | **0.045** | **0.063** |
| Tiredness<br>(frequency) | **-0.034** | 0.024 | 0.027 | **0.038** | -0.009 | -0.008 | **-0.027** | 0.025 | 0.009 | -0.011 | 0.023 | 0.008 | **0.096** |
| Headaches<br>(frequency) | **-0.036** | -0.009 | 0.023 | 0.020 | 0.017 | **0.032** | -0.002 | 0.010 | 0.010 | -0.011 | 0.015 | 0.008 | **0.068** |
| Reproductive/<br>mating success: |  |  |  |  |  |  |  |  |  |  |  |  |  |
| Number of<br>children | **0.023** | -0.007 | -0.026 | **-0.035** | 0.018 | 0.014 | -0.006 | -0.025 | -0.007 | **0.039** | **-0.032** | **0.099** | **-0.030** |
| Number of<br>sexual<br>partners | 0.018 | -0.002 | 0.020 | 0.025 | **-0.046** | 0.000 | **-0.019** | -0.005 | -0.003 | 0.009 | 0.009 | **0.069** | **0.182** |
| Negative health<br>effects | 1 | 0 | 2 | 6 | 9 | 3 | 3 | 1 | 0 | 3 | 11 | 10 | 10 |
| Positive health<br>effects | 9 | 2 | 0 | 1 | 1 | 2 | 13 | 0 | 0 | 3 | 0 | 3 | 10 |

*The figures (age-controlled partial Kendall Tau correlations) show the strength and direction of associations between variables on the top and on the left. A positive figure means a positive association between a respondent characteristic (column headings) and a category of human health, including number of children and sexual partners (row headings). Associations that remain significant after correction for multiple testing are in bold. The last two rows show the total number of significant associations where the effect on health is either negative or positive. A higher number of children and a higher number of sexual partners are classified as positive health effects.*

**Table 4.** Results for women: associations with eye color, hair color, hair waviness, BMI, and smoking

|  | Blue | Green | Brown | Black | Gray | Amber | Hazel | Eye<br>darkness | Hair<br>darkness | Hair<br>redness | Hair<br>waviness | BMI | Smoking |
| --- | --- | --- | --- | --- | --- | --- | --- | --- | --- | --- | --- | --- | --- |
| Physical health<br>problems in<br>general | -0.007 | **-0.024** | **0.039** | **-0.027** | 0.010 | -0.020 | -0.009 | 0.011 | **0.020** | **0.020** | -0.001 | **0.228** | **0.054** |
| Mental health<br>problems in<br>general | **-0.033** | 0.014 | 0.013 | **-0.024** | **0.019** | 0.001 | -0.007 | 0.003 | **0.020** | **0.047** | 0.008 | 0.013 | **0.041** |
| Specific health<br>problems: |  |  |  |  |  |  |  |  |  |  |  |  |  |
| Antibiotics/<br>year | -0.005 | **-0.026** | **0.020** | 0.012 | 0.015 | -0.005 | -0.001 | 0.002 | 0.013 | **-0.035** | 0.010 | **0.045** | **0.027** |
| Acute care/<br>5 years | **-0.025** | -0.013 | **0.035** | 0.015 | 0.010 | -0.007 | -0.009 | **0.019** | **0.025** | 0.002 | **0.018** | **0.047** | **0.029** |
| Different med.<br>specialist seen<br>every 2 years | -0.018 | -0.014 | **0.046** | 0.006 | -0.005 | -0.022 | -0.014 | **0.023** | 0.014 | -0.006 | 0.013 | **0.052** | 0.013 |
| Number of<br>prescription<br>drugs/day | **-0.055** | **-0.023** | **0.077** | -0.009 | **0.021** | 0.002 | **-0.028** | **0.045** | **0.023** | 0.008 | -0.013 | **0.122** | 0.005 |
| Number of<br>alternative<br>medicines/day | 0.001 | 0.012 | 0.003 | 0.008 | -0.012 | -0.022 | 0.001 | 0.007 | 0.004 | 0.013 | 0.012 | **-0.019** | -0.019 |
| Allergies | **-0.039** | -0.001 | **0.049** | 0.011 | -0.011 | -0.001 | -0.002 | **0.034** | 0.016 | -0.012 | **0.025** | **0.030** | **-0.025** |
| Immunological | **-0.020** | -0.015 | **0.034** | **0.040** | **-0.020** | 0.019 | 0.006 | **0.039** | **0.039** | 0.007 | **0.018** | **0.020** | **0.018** |
| Digestive | -0.015 | 0.018 | 0.016 | 0.012 | -0.002 | 0.010 | **-0.037** | 0.014 | -0.001 | 0.008 | **0.025** | **0.031** | 0.009 |
| Heart &<br>vascular<br>system | -0.010 | 0.009 | 0.005 | -0.010 | 0.011 | -0.006 | **-0.019** | 0.002 | 0.015 | **0.044** | **0.021** | **0.040** | **0.017** |
| Hematological | 0.004 | -0.016 | 0.000 | **0.036** | 0.008 | -0.014 | 0.009 | -0.004 | 0.000 | 0.007 | **0.039** | -0.006 | -0.007 |
| Metabolic | **-0.041** | **-0.022** | **0.039** | 0.019 | 0.010 | 0.021 | 0.014 | **0.038** | **0.023** | **0.045** | **0.020** | **0.187** | 0.007 |
| Cancer | 0.012 | **0.024** | -0.014 | -0.012 | **-0.027** | 0.017 | -0.010 | 0.003 | -0.009 | **0.067** | 0.013 | 0.011 | 0.013 |
| Fertility | **-0.030** | 0.005 | 0.015 | -0.005 | 0.013 | 0.007 | 0.001 | 0.013 | 0.015 | **0.042** | **0.054** | 0.006 | -0.010 |
| Genitourinary | -0.017 | -0.007 | **0.033** | -0.006 | **0.019** | -0.024 | **-0.025** | 0.016 | 0.009 | **0.028** | -0.011 | **-0.026** | 0.013 |
| Sense organs | -0.002 | -0.007 | -0.016 | **-0.034** | **0.038** | 0.005 | 0.003 | **-0.029** | 0.000 | 0.011 | **0.033** | **0.041** | **-0.024** |
| Neurological | -0.013 | -0.003 | 0.006 | **0.023** | **0.021** | -0.001 | **-0.020** | -0.002 | 0.011 | 0.014 | 0.015 | 0.011 | -0.005 |
| Psychiatric | **-0.033** | **0.025** | **-0.020** | -0.012 | **0.035** | 0.004 | 0.005 | -0.006 | 0.006 | **0.020** | 0.009 | 0.007 | **0.098** |
| Sexual<br>function | -0.012 | 0.003 | 0.012 | -0.016 | 0.016 | 0.006 | **-0.024** | -0.002 | 0.015 | **0.030** | 0.013 | 0.017 | **0.016** |
| Musculoskeletal | 0.000 | 0.010 | -0.005 | -0.007 | 0.005 | -0.015 | -0.005 | -0.004 | -0.018 | **0.053** | 0.007 | **0.040** | 0.013 |
| Respiratory | **-0.026** | 0.006 | **0.027** | **0.026** | -0.006 | -0.008 | -0.008 | **0.026** | 0.013 | -0.006 | 0.011 | **0.084** | **0.053** |
| Tiredness<br>(frequency) | -0.004 | 0.005 | 0.001 | 0.003 | 0.016 | -0.007 | **-0.022** | -0.005 | 0.014 | 0.014 | 0.002 | **0.021** | **0.058** |
| Headaches<br>(frequency) | -0.001 | 0.006 | -0.004 | 0.013 | 0.010 | -0.009 | **-0.019** | -0.009 | 0.015 | -0.002 | **0.018** | 0.014 | **0.019** |
| Reproductive/<br>mating success: |  |  |  |  |  |  |  |  |  |  |  |  |  |
| Number of | 0.006 | 0.004 | **0.026** | -0.008 | -0.016 | -0.010 | **-0.041** | **0.018** | **0.030** | **0.143** | **-0.026** | **0.096** | **-0.027** |
| children |  |  |  |  |  |  |  |  |  |  |  |  |  |
| Number of sexual partners | 0.010 | 0.016 | -0.006 | 0.014 | **-0.027** | 0.014 | -0.008 | -0.001 | **-0.021** | **0.061** | -0.008 | 0.011 | **0.246** |
| Negative health effects | 0 | 2 | 10 | 4 | 7 | 0 | 1 | 7 | 7 | 10 | 11 | 14 | 12 |
| Positive health effects | 9 | 4 | 2 | 3 | 2 | 0 | 8 | 2 | 1 | 3 | 0 | 3 | 3 |

*See Table 3 legend.*

The results shown in Table 3 and Table 4 suggest that women have more negative health effects associated with hair or eye color. Red hair in particular seems associated in women with the most negative effects and the fewest positive effects. To measure the size of health effects that disproportionately affect red-haired women, we performed MANCOVAs (multivariate analyses of variance) on the ones found to be significant. These effect sizes were compared with those of the three benchmarks: hair waviness, BMI, and smoking. By order of importance, red-haired women were prone to disorders in the following health categories: (1) Musculoskeletal; (2) Heart & vascular, Cancer, Fertility; (3) Metabolic; (4) Sexual function; (5) Genitourinary. The sizes of these health effects seemed comparable to those of BMI and smoking, though smaller.

To determine the relative importance of skin cancer in the Cancer category, we looked at the incidence of specific disorders within that category. The results are shown in Table 5.

**Table 5.** Differences in cancer rate between redheads and non-redheads by sex and by specific type of cancer

| Type of cancer | Men |  |  |  |  |  | Women |  |  |  |  |  |
| --- | --- | --- | --- | --- | --- | --- | --- | --- | --- | --- | --- | --- |
|  | Red-<br>Can- | Red-<br>Can+ | Red+<br>Can- | Red+<br>Can+ | OR | p | Red-<br>Can- | Red-<br>Can+ | Red+<br>Can- | Red+<br>Can+ | OR | p |
| Esophageal cancer | 1762 | 0 | 268 | 0 |  |  | 2946 | 1 | 836 | 1 | 3.52 | 0.394 |
| Stomach cancer | 1762 | 0 | 268 | 0 |  |  | 2946 | 1 | 836 | 1 | 3.52 | 0.394 |
| Colorectal cancer | 1762 | 0 | 264 | 4 | 273.56 | **0.000** | 2947 | 0 | 834 | 3 | 109.53 | **0.011** |
| Liver cancer | 1762 | 0 | 267 | 1 | 72.57 | 0.132 | 2947 | 0 | 836 | 1 | 38.77 | 0.221 |
| Lung cancer | 1761 | 1 | 267 | 1 | 6.59 | 0.247 | 2947 | 0 | 837 | 0 |  |  |
| Melanoma, other skin cancers | 1753 | 9 | 266 | 2 | 1.52 | 0.271 | 2937 | 10 | 833 | 4 | 1.43 | 0.243 |
| Breast cancer | 1762 | 0 | 268 | 0 |  |  | 2932 | 15 | 830 | 7 | 1.66 | 0.318 |
| Cervical uterine precancerosis | 1677 | 0 | 259 | 0 |  |  | 2723 | 64 | 756 | 27 | 1.52 | **0.008** |
| Cervical uterine cancer | 1762 | 0 | 268 | 0 |  |  | 2902 | 45 | 817 | 20 | 1.58 | **0.003** |
| Corpus uteri cancer | 1762 | 0 | 268 | 0 |  |  | 2940 | 7 | 835 | 2 | 1.04 | 1.000 |
| Ovarian cancer | 1762 | 0 | 268 | 0 |  |  | 2942 | 5 | 832 | 5 | 3.54 | **0.015** |
| Prostate cancer | 1753 | 9 | 266 | 2 | 1.52 | 0.184 | 2947 | 0 | 837 | 0 |  |  |
| Lymphoma, myeloma<br>multiple | 1760 | 2 | 268 | 0 | 0.31 | 1.000 | 2947 | 0 | 837 | 0 |  |  |
| Leukemia | 1758 | 4 | 268 | 0 | 0.16 | 1.000 | 2941 | 6 | 836 | 1 | 0.63 | 1.000 |
| Bladder cancer | 1761 | 1 | 268 | 0 | 0.60 | 1.000 | 2945 | 2 | 836 | 1 | 1.85 | 0.528 |
| Mouth, oropharynx cancers | 1761 | 1 | 268 | 0 | 0.60 | 1.000 | 2947 | 0 | 836 | 1 | 38.77 | 0.221 |
| Adeno-carcinoma | 1762 | 0 | 268 | 0 |  |  | 2943 | 4 | 837 | 0 | 0.09 | 0.582 |
| Papilloma cancer | 1762 | 0 | 268 | 0 |  |  | 2941 | 6 | 833 | 4 | 2.37 | 0.422 |
| Other types of cancer | 1747 | 15 | 265 | 3 | 1.35 | 0.642 | 2929 | 18 | 825 | 12 | 2.37 | **0.000** |

*Red- Can- = Number of non-redheads (i.e., respondents whose intensity of redness is 1-3 on a 6-point Likert scale) without the specific type of cancer*

*Red- Can+ = Number of non-redheads with the specific type of cancer*

*Red+ Can- = Number of redheads (i.e., respondents whose intensity of redness is 4-6) without the specific type of cancer*

*Red+ Can+ = Number of redheads with the specific type of cancer*

*Odds Ratios (OR) and statistical significance (p) respectively are shown for men and women and for each specific type of cancer. The effect of age on health status was controlled by performing partial Kendall’s correlation whenever the incidence of a specific type of cancer exceeded 9. Otherwise, Fisher’s exact test was performed to determine statistical significance. ORs higher than 1 indicate that redness is positively associated with the incidence of the specific type of cancer. Results are in bold if significant in two-sided tests after Benjamini-Hochberg correction for multiple tests, and p-values < 0.0005 are coded as 0.000.*

Red-haired men were thus more prone to colorectal cancer, while red-haired women were more prone to colorectal cancer, precancerous cervical or uterine lesions, cervical or uterine cancer, ovarian cancer, and other types of cancer. This higher cancer risk was not due to a higher rate of skin cancer, which was only non-significantly more frequent in red-haired men (OR=1.52) and women (OR=1.43).

### Gender effects and interactions

Because health effects differed between men and women, particularly among red-haired respondents, we investigated whether gender interacted significantly with the apparent effects of hair/eye color on their health. To this end, we first performed four MANCOVAs to see whether variance in respondent health correlated significantly with gender, eye darkness, hair darkness, and hair redness. We then performed three MANCOVAs to see whether variance in respondent health correlated significantly with an interaction between gender and any of the other variables: eye darkness, hair darkness, or hair redness. Finally, we constructed a new binary variable—presence or absence of green eyes—and performed two MANCOVAs to see whether variance in respondent health correlated significantly with this new variable or with an interaction between it and gender. The results are shown in Table 6.

**Table 6.** Correlations of all health effects with gender, hair color, or eye color

| Independent variable | R | p-value |
| --- | --- | --- |
| gender | 0.12 | 0.001*** |
| hair redness | 0.04 | 0.001*** |
| hair darkness | 0.02 | 0.35 |
| eye darkness | 0.03 | 0.002** |
| gender*hair redness | 0.02 | 0.182 |
| gender*hair darkness | 0.01 | 0.583 |
| gender*eye darkness | 0.02 | 0.405 |
| green eyes vs. all other eye colors | 0.02 | 0.049* |
| gender*green eyes vs. all other eye colors | 0.12 | 0.001*** |

Although respondent health correlated significantly with gender, hair redness, and eye darkness taken separately, and although female respondents differed significantly from male respondents in hair redness and hair darkness (but not eye darkness), there were no significant three-way interactions between gender, respondent health, and any of the above variables for hair or eye color (Table 6). In the case of eye color, a linear regression on eye darkness may not be the best way to capture a combined effect by gender and eye color on respondent health. Indeed, the relationship between gender and eye color cannot be described simply in terms of eye darkness. As we have seen, women are less likely than men to be blue-eyed or brown-eyed, while conversely being more often green-eyed (Table 1). This was why we constructed the binary variable of green eyes versus all other eye colors, and we found that this variable significantly interacted with gender to produce effects on respondent health. In general, green-eyed women were healthier than the other respondents, except for a greater propensity to have cancer and psychiatric problems.

This finding made us take a second look at the relationship between female respondent health and hair redness. That relationship, too, might not be fully understood through a linear regression. We specifically looked at the data on cancer because the relationship between health outcomes and hair redness was strongest in that category, even though the well-known association between red hair and skin cancer seemed to contribute very little to this relationship. We performed a logistic regression with the incidence of any cancer as the dependent variable (0 = no cancer reported, 1 = cancer or precancerous lesion reported) and with three independent variables: gender, age, hair redness, and gender*hair redness interaction. A separate analysis for women showed that hair redness (OR range = 3.99, p<0.0001) and age (OR range = 12.1, p<0.0001) significantly affected the incidence of any cancer. In contrast, a separate analysis for men showed a significant age effect (OR range = 50.7, p<0.0001) but no hair redness effect (OR range = 1.63, p=0.457). For men and women together, cancer was significantly affected by age (OR range = 17.7, p<0.0001) and by hair redness (OR range = 3.74, p<0.0001) but not by gender (OR range = 0.67, p=0.331) or by gender*hair redness (OR range = 0.47, p=0.381). The results are shown in Figure 5.

**Figure 5.**
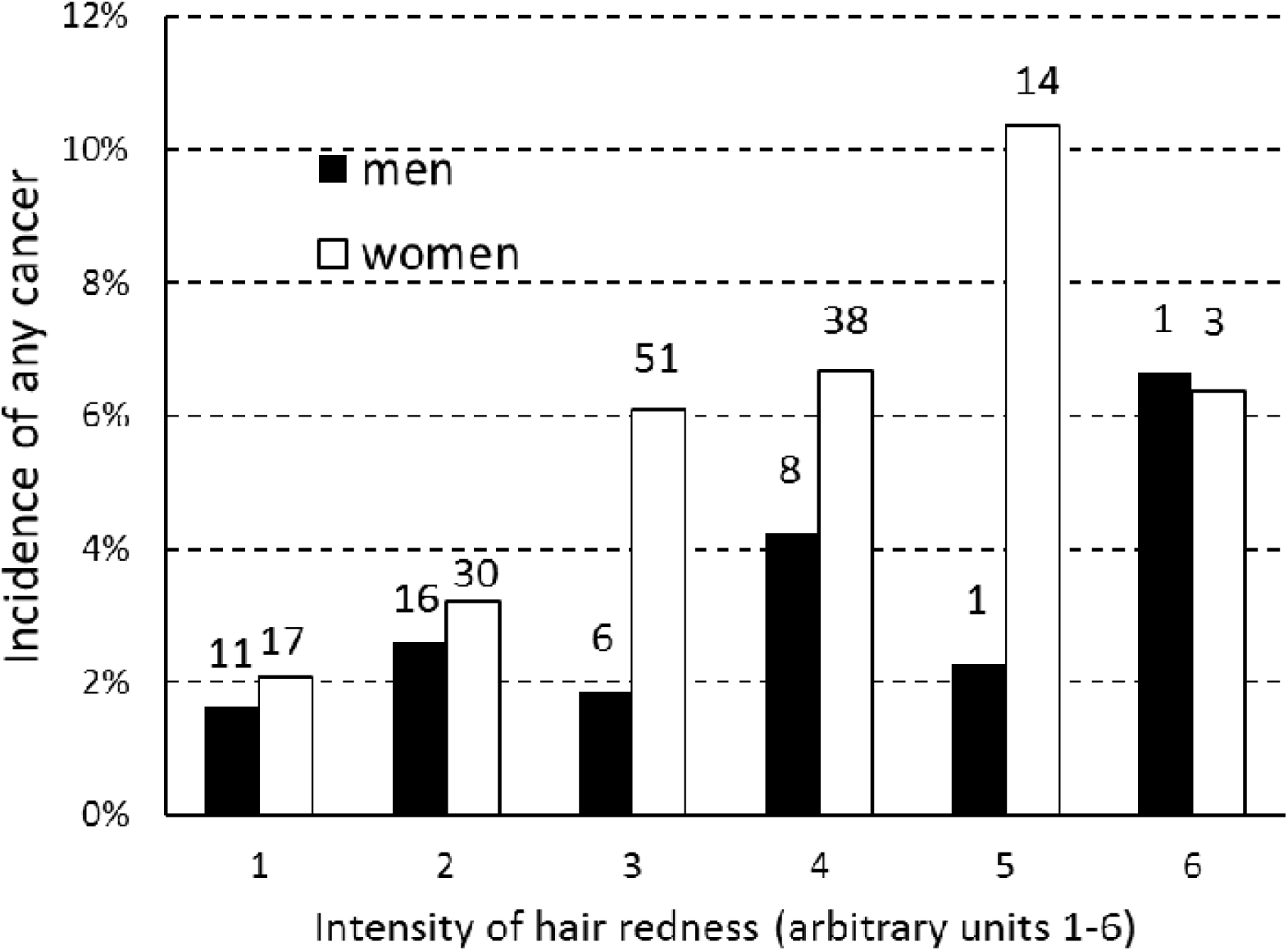
Incidence of any cancer by gradation of hair redness, for men and women

*The numbers above the columns show numbers of subjects in particular categories.*

The gender difference was greatest at the next-to-last gradation of hair redness. To learn more about this interaction between gender and gradation of hair redness, we plotted the reported mean seriousness o cancer (where 1 = no cancer reported and 6 = very serious problem with cancer) as a function of hair redness. This new variable may provide a clearer picture because it contains more information than simply the presence or absence of cancer. The results are shown in Figure 6. Mean seriousness of cancer increased steadily with increasing hair redness up to a certain gradation of redness and then decreased. This relationship was stronger in women than in men and peaked at a higher gradation of redness in women th in men. For women, both age (p<0.0005) and hair redness (p=0.014) significantly affected mean seriousness of cancer. Men showed significant effects for age (p<0.0005) but not for hair redness (p=0.719). For men and women together, mean seriousness of cancer was significantly affected by age (p<0.005) and by hair redness (p=0.0005) but not by gender (p=0.438) or by gender*hair redness (p=0.292).

**Figure 6.**
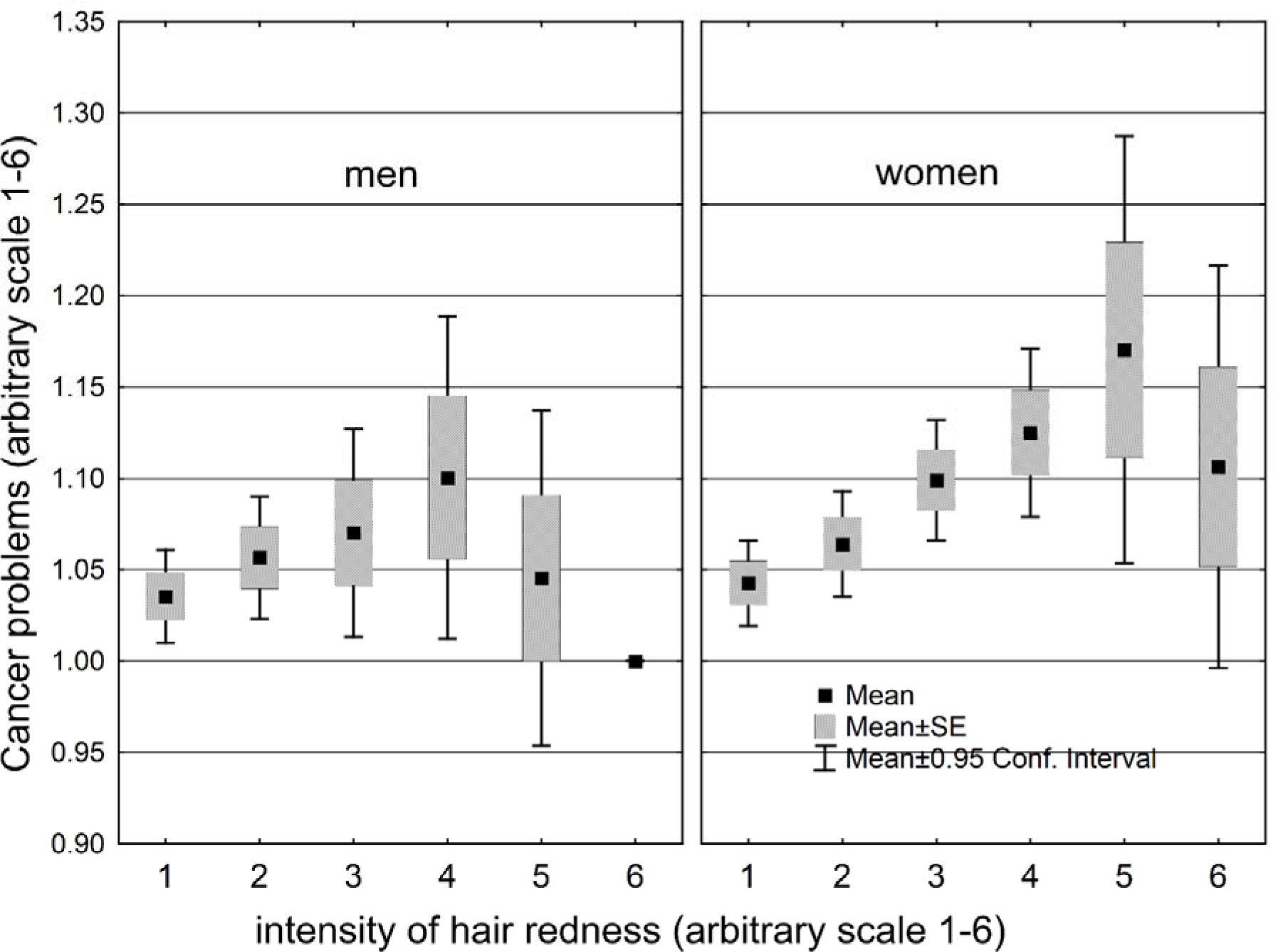
Mean seriousness of cancer by gradation of hair redness, for men and women

## Discussion

Red hair seems to be costly for women’s health. In this study, red-haired women did worse than other women in ten health categories and better in only three. In general, women incurred more costs and gained fewer benefits from red hair than from any other hair or eye color. Brown eyes held second place, but the health effects associated with brown eyes, both negative and positive, were smaller on average than those associated with red hair. Red-haired men showed a balanced pattern of health effects, doing better than other men in three categories and worse in three. Number of children was the only category where both male and female redheads did better than non-redheads. In terms of reproductive and, ultimately, evolutionary success, red hair seems to be a plus rather than a minus.

The cancer rate was higher among red-haired women than among other women, and we initially suspected a higher rate of skin cancer as the cause. A closer look at the data, however, showed that the higher cancer rate was due not to a higher incidence of skin cancer, but rather to a higher incidence of cancers in the colorectal region, the cervix, the uterus, and the ovaries (Table 5). Because estrogen influences the development of the last three organs from the fetal stage onward, the higher cancer rate may be better explained by a higher level of prenatal exposure to estrogen, rather than by greater vulnerability to UV. This explanation is supported by the higher incidence among red-haired women of osteoporosis and obstetric complications (results not shown), both of which are either more frequent in women or specific to women. It may seem surprising that the skin cancer rate was only slightly higher for redheads than for non-redheads, given the many studies that point to red hair as a risk factor. Such studies, however, generally concern countries like the United States and Australia, whose citizens are exposed to a higher intensity of UV because they live at lower latitudes than do Czech citizens and also because a higher proportion of them have been regularly traveling to tropical or subtropical resorts for the past half-century or longer.

If we consider the other negative health effects associated with red hair, these too are not easily attributable to fairness of skin, and hence to UV vulnerability, again because of the greater propensity of women to exhibit these health effects. Although women are fairer-skinned than men, this gender difference is smaller in fair-skinned humans and in redheads in particular, among whom both sexes are pushed up against the physiological “ceiling” of skin reflectance [23, 24]. Moreover, if vulnerability to UV explains this pattern of health effects, we would expect to see a similar pattern with blue eyes, which are likewise associated with fair skin [1]. Yet, relative to other hair and eye colors in our sample, blue eyes imposed fewer costs on male or female health, while providing women with the highest total of benefits and men the second-highest.

Some of these other negative health effects are consistent with previous findings in the literature. Despite having more children on average, the red-haired women of this study had a higher incidence of fertility problems, which would be consistent with the higher incidence of endometriosis reported in previous studies. They also had more neurological problems, although none of these involved Parkinson’s disease. Actually, few cases of Parkinson’s would be expected, given the relatively young age of the respondents. Red-haired women showed no obvious indications of increased pain sensitivity in this study, although in some cases they might have reported more medical problems because sensitivity to pain made them seek medical assistance more readily.

These health effects thus seem to be due to a female-specific factor that is most strongly expressed in red-haired women. The relationship between this factor and hair redness seems curvilinear, i.e., average female health progressively worsens with redder gradations of hair, but only up to a certain point. If we take the data on seriousness of cancer, the worst health was reported by women with the next-to-last gradation of hair redness. Those with the reddest hair were actually somewhat better off (Figure 5); however, this category had only three women and one man. With respect to eye color, the female-specific factor seemed to act most strongly on women with green eyes and not on those with lighter shades. In both cases, these effects mirror the effects of estrogen on development of hair and eye color. Red hair is more frequent in women than in men (Figure 1); similarly, green eyes are more frequent in women than in men, with brown eyes and blue eyes showing the reverse pattern (Table 1).

In addition to the problem of modeling the curvilinear relationships between female respondent health and eye darkness/hair redness, the data suffered from a high level of noise. Inter-respondent variance was inflated by self-report and the subjective nature of the questions. As a result, even when significant correlations were found between respondent health and different variables (gender, age, hair redness, hair darkness, eye darkness, etc.), they cannot explain more than a tiny proportion of total variance in health status among the respondents. We should emphasize that this tininess may be only apparent. To provide a benchmark for the relative importance of these health effects, we also examined respondent data on BMI and smoking, both of which strongly affect human health. Using the data in Table 4 (significant negative effects on women’s health), we found that, on average, BMI explains twice as much variance in these negative health effects as does red hair (Tau=0.071, vs. 0.040), while smoking explains approximately the same amount (Tau=0.039). Thus, BMI and smoking overlap with hair and eye color in the magnitude of their negative effects on women’s health.

It seems, then, that different hair and eye colors are associated with significantly different health outcomes and that these apparent effects are stronger in women than in men. Red-haired women exhibit the most divergent health effects, including a previously unreported vulnerability to colorectal, ovarian, and cervical cancer. Not all effects are for the worse. Red-haired women seem to enjoy greater reproductive and mating success, as measured by number of children and number of sexual partners. It may be that red-haired women have more children because they begin having them at an earlier age, although a recent study has reported that red-haired men and women lose their virginity at a later age on average [25]. An alternative explanation is that red-haired women attract not only more sexual partners but also better sexual partners who can support a larger family size. More research on the life history of red-haired women is needed. Red hair might be more attractive than other hair colors because it is less common. It has been argued that the different hair and eye colors of Europeans, including red hair, have coexisted in a dynamic equilibrium [13,14]. According to this argument, a hair or eye color is sexually attractive in proportion to its scarcity. It therefore loses this novelty value if it becomes too common, and the pressure of selection shifts to less common variants. In the case of red hair, there may also be an equilibrium between sexual attractiveness and negative health effects. These negative effects would depress the population frequency of red hair below the frequency it would have if sexual attractiveness were the only selection pressure. The gap between this second equilibrium and the first may explain the relative popularity of red-haired women: they have never been sufficiently numerous to lose their novelty value.

What causes these negative health effects in red-haired women? The cause can be framed in either biochemical or evolutionary terms. First, these effects might be inherent to biosynthesis of red hair pigments (pheomelanins). But why, then, are they expressed much more often in red-haired women than in red-haired men? There seems to be a female-specific factor. As argued in the Introduction, this factor may be prenatal estrogen. The same prenatal estrogen that causes red hair to be more frequent in women than in men may also explain why these negative health effects are expressed in red-haired women but not in red-haired men. In the fetal stage, such women were more likely to experience estrogen levels near the top end of the normal range for fetal development. The risk of later health problems would therefore be proportionately greater.

Second, in terms of evolutionary causation, red hair may have been the last hair color to emerge in modern humans; therefore, not enough time has passed for corrective evolution, either through new alleles that produce red hair with fewer side effects or through modifier genes that neutralize the side effects of existing red hair alleles. This situation is typical of rapid evolution over relatively short spans of time [26]. Another possible scenario is that red hair alleles emerged among Neanderthals and then introgressed into early modern Europeans when the two groups coexisted in Europe. Such introgression could cause genetic incompatibility and thus explain the negative health effects we observed. Red hair is produced mainly by five loss-of-function alleles at the *MCIR* gene, and a recent study has identified one of them, Val92Met, as a likely Neanderthal introgression. The same study, however, found that the four other alleles are not of Neanderthal origin [27]. Given that modern humans entered Europe c. 45,000 BP and reached northern Europe c. 30,000 BP, and that the Neanderthals went extinct c. 40,000 BP, the scenario of Neanderthal introgression makes sense for Val92Met, which is found throughout Eurasia. However, the other loss-of-function alleles, which are more specific to northern Europe, probably originated in modern humans.

To conclude, our findings may shed light not only on the health risks associated with red hair but also on the evolution of this highly visible color trait and, more generally, on how the diverse European palette of hair and eye colors came into being. This evolution seems to have occurred for the most part in relatively recent times, probably no earlier than the entry of modern humans into northern Europe some 30,000 years ago and no later than the oldest DNA evidence of these alternate colors, dating to some 8,000 years ago from Motala, Sweden. The short time span (< 1000 generations) suggests that some form of selection, possibly sexual selection, was driving this diversification of hair and eye colors in early Europeans. Of these ‘new’ color variants, red hair seems to diverge the most from the ancestral state of black hair and brown eyes. It is the most sexually dimorphic variant, not only in population frequency but also in health outcomes.

## Declarations

### Acknowledgements and Funding

The work was supported by Charles University (UNCE 204004) and the Czech Science Foundation (Grant No. P407/16/20958).

## References

1. Scherer D, Kumar R. Genetics of pigmentation in skin cancer - a review, Mutat Res. 2010; 705: 141-153.

2. Andresen T, Lunden D, Drewes AM, Arendt-Nielsen L. Pain sensitivity and experimentally induced sensitisation in red haired females, Scand J Pain. 2011; 2: 3-6.

3. Gao X, Simon KC, Han J, Schwarzchild MA, Ascherio A. Genetic determinants of hair color and Parkinson's disease risk, Ann Neurol. 2009; 65: 76-82.

4. Liem EB, Joiner TV, Tsueda K, Sessler DI. Increased sensitivity to thermal pain and reduced subcutaneous lidocaine efficacy in redheads, Anesthesiology. 2005; 102: 509-514.

5. Somigliana E, Viganò P, Abbiati A, Gentilini D, Parazzini F, Benaglia L, Vercellini P, Fedele L. ‘Here comes the sun’: pigmentary traits and sun habits in women with endometriosis Hum Reprod. 2010; 25: 728-773.

6. Tell-Marti G, Puig-Butille JA, Potrony M, Badenas C, Mila M, Malvehy J, Marti MJ, Ezquerra M, Fernandez-Santiago R, Puig S. The *MC1R* melanoma risk variant p.R160W is associated with Parkinson disease, Ann Neurol. 2015; 77: 889-894.

7. Woodworth SH, Singh M, Yussman MA, Sanfilippo JS, Cook CL, Lincoln SR. A prospective study on the association between red hair color and endomtriosis in infertile patients, Fertil Steril. 1995; 64: 651-652.

8. Chen X, Chen H, Cai W, Maguire M, Ya B, Zuo F, Logan R, Li H, Robinson K, Vanderburg CR, Yu Y, Wang Y, Fisher DE, Schwarzschild, MA. The melanoma-linked "redhead" *MC1R* influences dopaminergic neuron survival, Ann Neurol. 2017; 81: 395-406.

9. Shekar SN, Duffy DL, Frudakis T, Montgomery GW, James MR, Sturm RA, Martin NG. Spectrophotometric methods for quantifying pigmentation in human hair—Influence of MC1R genotype and environment, Photochem Photobiol. 2008; 84: 719-726.

10. Kleisner K, Kočnar T, Rubešová A, Flegr J. Eye color predicts but does not directly influence perceived dominance in men, Pers Indiv Differ. 2010; 49: 59-64.

11. Kleisner K, Priplatova L, Frost P, Flegr J. Trustworthy-Looking Face Meets Brown Eyes. PLoS ONE. 2013; 8, e53285.

12. Martinez-Cadenas C, Peña-Chilet M, Ibarrola-Villava M, Ribas G. Gender is a major factor explaining discrepancies in eye colour prediction based on *HERC2/OCA2* genotype and the IrisPlex model. Forensic Sci Int-Gen. 2013; 7:453-60.

13. Frost P. European hair and eye color. A case of frequency-dependent sexual selection? Evol Hum Behav. 2006; 27: 85-103.

14. Frost P. The Puzzle of European Hair, Eye, and Skin Color. Adv Anthropol. 2014; 4: 78-88.

15. Kankova S, Flegr J, Calda P. The influence of latent toxoplasmosis on women's reproductive function: four cross-sectional studies. Folia Parasitol. 2015; 62.

16. Flegr J, Hoffmann R, Dammann M. Worse health status and higher incidence of health disorders in Rhesus negative subjects. PLoS ONE. 2015; 10.

17. Flegr J, Hodny Z. Cat scratches, not bites, are associated with unipolar depression - cross-sectional study. Parasite Vector. 2016; 9.

18. Oksanen J, Blanchet FG, Friendly M, Kindt R, Legendre P, McGlinn D, Minchin PR, O'Hara RB, Simpson GL, Solymos P, Stevens HH, Szoecs E, Wagner E vegan: Community Ecology Package. R package version 2.4-1., 2016; https://CRAN.R-project.org/package=vegan

19. Kaňková Š, Kodym P, Flegr J. Direct evidence of *Toxoplasma*-induced changes in serum testosterone in mice. Exp Parasitol. 2011; 128:181-3.

20. Siegel S, Castellan NJ. Nonparametric statistics for the behavioral sciences. 2nd ed. New York: McGraw-Hill; 1988.

21. Benjamini Y, Hochberg Y. Controlling the false discovery rate: A practical and powerful approach to multiple testing. J Roy Stat Soc B Met. 1995; 57:289-300.

22. Simpson EH. Measurement of Diversity, Nature. 1949; 163:688

23. Frost P. Comment on Human skin-color sexual dimorphism: A test of the sexual selection hypothesis, Am J Phys Anthropol. 2007; 133: 779-781.

24. Madrigal L, Kelly W. Human skin-color sexual dimorphism: a test of the sexual selection hypothesis, Am J Phys Anthropol. 2006; 132: 470-482.

25. Day FR, Helgason H, Chasman DI, Rose LM, Loh P-R, Scott RA, Helgason A, Kong A, Masson G, Magnusson OT, Gudbjartsson D, Thorsteinsdottir U, Buring JE, Ridker PM, Sulem P, Stefansson K, Ong KK, Perry JRB. Physical and neurobehavioral determinants of reproductive onset and success, Nat Genet. 2016; 48: 617-623.

26. Flegr J. Elastic, not plastic species: Frozen plasticity theory and the origin of adaptive evolution in sexually reproducing organisms, Biol Direct. 2010; 5:2.

27. Ding Q, Hu Y, Xu S, Wang C-C, Li H, Zhang R, Yan S, Wang J, Jin L. Neanderthal origin of the haplotypes carrying the functional variant Val92Met in the *MC1R* in modern humans, Mol Biol Evol. 2014; 31:1994-2003.

